# Assessing auditory processing endophenotypes associated with Schizophrenia in individuals with 22q11.2 Deletion Syndrome

**DOI:** 10.1101/696021

**Authors:** Ana A. Francisco, John J. Foxe, Douwe J. Horsthuis, Danielle DeMaio, Sophie Molholm

## Abstract

**Background:** 22q11.2 Deletion Syndrome (22q11.2DS) is the strongest known molecular risk factor for schizophrenia. Brain responses to auditory stimuli have been studied extensively in schizophrenia and described as potential biomarkers of vulnerability to psychosis. We sought to understand whether these responses might aid in differentiating individuals with 22q11.2DS as a function of psychotic symptoms, and ultimately serve as signals of risk for schizophrenia.

**Methods:** A duration oddball paradigm and high-density electrophysiology were used to test auditory processing in 26 individuals with 22q11.2DS (13-35 years old, 17 females) with varying degrees of psychotic symptomatology and in 26 age- and sex-matched neurotypical controls (NT). Presentation rate varied across three levels, to examine the effect of increasing demands on memory and the integrity of sensory adaptation. We tested whether N1 and mismatch negativity (MMN), typically reduced in schizophrenia, related to clinical/cognitive measures, and how they were affected by presentation rate.

**Results:** N1 adaptation effects interacted with psychotic symptomatology: Compared to an NT group, individuals with 22q11.2DS but no psychotic symptomatology presented larger adaptation effects, whereas those with psychotic symptomatology presented smaller effects. In contrast, individuals with 22q11.2DS showed increased effects of presentation rate on MMN amplitude, regardless of the presence of symptoms. While IQ and working memory were lower in the 22q11.2DS group, these measures did not correlate with the electrophysiological data.

**Conclusions:** These findings suggest the presence of two distinct mechanisms: One intrinsic to 22q11.2DS resulting in increased N1 and MMN responses; another related to psychosis leading to a decreased N1 response.

## Introduction

22q11.2 Deletion Syndrome (22q11.2DS henceforth; also named DiGeorge syndrome or velo-cardio-facial syndrome) results from a hemizygous microdeletion of approximately 1.5 to 3 megabases on the long arm of chromosome 22. The deleted region contains about 60 known genes, some of which are highly expressed in the brain and known to affect early neuronal migration and cortical development (1, 2). 22q11.2DS is the most common chromosomal microdeletion disorder, with prevalence estimates ranging from 1 per 1000 to 1 per 4000 live births (3, 4).

The phenotypic expression of 22q11.2DS is highly variable and ranges from life-threatening to less severe conditions (5). Its clinical presentation includes variable developmental delays, cognitive deficits and neuropsychiatric conditions, and multi-organ dysfunction such as cardiac and palatal abnormalities (6, 7). Cognitively, 22q11.2DS is characterized by deficits in executive function (8–11), nonverbal memory (12, 13), visuospatial (14–16) and visual-motor (17) processing, and working memory (18). Approximately 60% of individuals diagnosed with 22q11.2DS meet criteria for at least one psychiatric diagnosis (19–21) and the development of psychosis is one of the most significant concerns for parents of children with 22q11.2DS: 20 to 40% of individuals identified with the deletion go on to develop schizophrenia (22–25).

Idiopathic and 22q11.2DS-associated schizophrenia present a similar clinical path leading to psychosis (26, 27), with subtle social deficits and poor neurocognitive performance preceding its onset both in the general population (28, 29) and in 22q11.2DS (30). The clinical presentation of schizophrenia does not appear to differ either (31), with deficits in cognition and perception, and positive (hallucinations, delusions, thought disorder) and negative (flat affect, alogia, avolition and anhedonia) symptoms (32). However, neurocognitive features are more severe in 22q11.2DS-associated schizophrenia, likely reflecting poorer baseline cognitive function (33). These similarities are accompanied by high concordance of neuroanatomic correlates (34–39), suggesting that common cerebral alterations may underlie psychotic symptomatology in both populations.

Reducing the duration of untreated psychosis, before irreversible pathological brain changes take place (40), leads to better functional outcomes in patients with psychotic disorders such as schizophrenia (41, 42), strongly motivating the goal of early identification prior to frank disease onset. Since clearly observable behavioral changes are often the product of processes that have started occurring in the brain significantly earlier, the direct and reliable measurement of functional brain activity and the consequent potential to detect neural vulnerability prior to behaviorally observable symptoms is key to enabling interventions focused on prevention rather than on treatment. Sensory and cognitive auditory evoked potentials (AEP) such as the N1 and the mismatch negativity (MMN) have been identified as potential biomarkers for vulnerability, conversion, and progression of psychosis in schizophrenia. The auditory N1 is the first prominent negative AEP (43), and reflects neural activity generated in and around primary auditory cortex (44). Several studies demonstrate a reduction of the auditory N1 in schizophrenia, in at-risk individuals and chronic and first-episode patients (45–49); but see (50) for a review on contradictory evidence. The MMN, in turn, operating at the sensory memory level, occurs when a repeating stimulus (the standard) in an auditory stream is replaced by a deviant stimulus: Regular aspects of consecutively presented standards form a memory trace; violation of those regularities by a deviant induces the MMN (51). Occurring typically 100 to 200 ms following the deviant event, the MMN is thought to reflect largely preattentive neural processes underlying detection of a pattern violation and updating of a representation of a regularity in the auditory environment (52–54). MMN amplitude has been shown to be reduced in schizophrenia (for reviews, see (55, 56)) in at-risk individuals (57–61), recent onset (58, 59, 62–64), and chronic patients (58, 62, 65–72), particularly when using duration deviants (65, 73, 74).

While previous studies have measured the N1 and MMN in 22q11.2DS, the limited results available have yielded inconsistent findings (75, 76). Such inconsistencies might reflect the relatively high degree of phenotypic variability that is inherent to this population. Here, we sought to measure the relationship between auditory brain function and cognitive function, focusing on cognitive abilities known to be vulnerable in this population. We also set out to test for potential associations between MMN and N1 amplitudes and the number of psychotic symptoms expressed in the 22q11.2DS cohort. Given the working memory difficulties previously described in 22q11.2DS, presentation rate (stimulus onset asynchronies: SOAs) was parametrically varied, which allowed us to examine not only the impact of increasing demands on the sensory memory system (77, 78), but also the integrity of sensory adaptation effects (77, 79).

## Methods and Materials

### Participants

Twenty-six participants diagnosed with 22q11.2DS (age range: 13-35 years old; 15 with at least one psychotic symptom) and 26 neurotypical age-matched controls (NT) (age range: 13-38 years old) were recruited. Exclusionary criteria for the neurotypical group included hearing impairment, developmental and/or educational difficulties or delays, neurological problems, and the presence of psychotic symptomatology. Exclusionary criteria for the 22q11.2DS group included hearing impairment and current neurological problems. Participants passed a hearing test (below 25dB HL for 500, 100, 2000, 4000Hz) performed on both ears using a *Beltone Audiometer* (Model 112). All participants signed an informed consent approved by the Institutional Review Board of the Albert Einstein College of Medicine and were monetarily compensated for their time. All aspects of the research conformed to the tenets of the Declaration of Helsinki.

### Experimental Procedure and Stimuli

Testing was carried out over a 2-day period and included cognitive testing, a diagnostic interview, and EEG recording. Cognitive testing focused on measures of intelligence, as assessed by the Wechsler Adult Intelligence Scale, WAIS-IV (80) or the Wechsler Intelligence Scale for Children, WISC-V (81). The IQ measure used refers to the Full-Scale IQ index. The working memory score used refers to the Working Memory Index, which comprises the Digit Span subtest and the Picture Span subtest (WISC-V) or the Arithmetic subtest (WAIS-IV). The Structured Clinical Interview for DSM-5, SCID-V (82) or the Structured Clinical Interview for DSM-IV Childhood Diagnoses, Kid-SCID (83) were performed to assess the presence of psychotic symptoms.

The EEG protocol focused on auditory processing, utilizing a traditional duration-MMN oddball paradigm. Participants sat in a sound- and electrically-shielded booth (Industrial Acoustics Company Inc, Bronx, NY) and watched a muted movie of their choice on a laptop (Dell Latitude E6430 ATG or E5420M) while passively listening to regularly (85%) occurring standard tones interspersed with deviant tones (15%). These tones had a frequency of 1000 Hz with a rise and fall time of 10 ms, and were presented at an intensity of 75dB SPL using a pair of Etymotic insert earphones (Etymotic Research, Inc., Elk Grove Village, IL, USA). Standard tones had a duration of 100 ms while deviant tones were 180 ms in duration. These tones were presented in a random oddball configuration (excepting that at least two standards preceded a deviant) to yield an MMN. In three blocked conditions, the stimulus onset asynchrony (SOA) was either 450 ms, 900 ms or 1800 ms. Each SOA was presented in blocks of 4 minutes long, composed of 500, 250 or 125 trials respectively. Participants were presented with 14 blocks (2*450 ms, 4*900 ms and 8*1800 ms) randomized across subjects, resulting in a possible 1000 trials (and 150 deviants) per SOA.

### Data acquisition and analysis

EEG data were acquired continuously at a sampling rate of 512 Hz from 71 locations using 64 scalp electrodes mounted on an elastic cap and seven external channels (mastoids, temples, and nasion) (Active 2 system; Biosemi^tm^, The Netherlands; 10-20 montage). Preprocessing was done using the EEGLAB (version 14.1.1) (84) toolbox for MATLAB (version 2017a; MathWorks, Natick, MA). Data were downsampled to 256 Hz, re-referenced to TP8 and filtered using a 1 Hz high pass filter (0.5 Hz transition bandwidth, filter order 1690) and a 45 Hz low pass filter (5 Hz transition bandwidth, filter order 152). Both were zero-phase Hamming windowed sinc FIR filters. Bad channels were automatically detected based on kurtosis measures and rejected after visual confirmation. Artifacts were removed by running an Independent Component Analysis (ICA) to exclude components accounting for eye blinks and saccades. After ICA, the previously excluded channels were interpolated, using the spherical spline method. Data were segmented into epochs of −100ms to 400ms using a baseline of −100ms to 0ms. These epochs went through an artifact detection algorithm (moving window peak-to-peak threshold at 120 µV). The number of trials included in the analyses did not differ between the groups (neurotypical control group: 597-1065 trials for standards, 96-193 for deviants; 22q11.2DS group: 578-956 trials for standards, 100-165 for deviants).

The definition of the N1 and the MMN windows was based on the typical time of occurrence of the N1 and duration-MMN components, and on visual confirmation that amplitudes were maximal in these intervals. N1 amplitude was measured between 80 and 120 ms on the standard waveform. The MMN is the difference between deviants and standards and was here measured between 200 and 240 ms (100 to 140 ms post deviance onset). Amplitude measures were taken at FCz, where signal was maximal for both groups. These amplitudes were used for between-groups statistics and correlations. All *p-values* (from *post-hoc* tests and correlations) were submitted to Holm-Bonferroni corrections for multiple comparisons (85), using the *p.adjust* of the *stats* package in R (86).

## Results

Table 1 shows a summary of the included participants’ IQ and performance on the working memory tasks. Working memory scores were only obtained for a subset of the individuals in the neurotypical group (N=18; the remainder were part of a control group for a study in which a reduced version of the IQ scale was used). Two-sample independent-means *t* tests were used to test for group differences. When the assumption of the homogeneity of variances was violated, *Welch* corrections were applied to adjust degrees of freedom. Statistical analyses confirmed that the groups differed significantly in IQ and working memory. There were significantly more females than males in each group, but no differences in sex between groups. Likewise, the groups did not differ significantly in age. 34.6% of those with 22q11.2DS were taking antidepressants, 23.1% anticonvulsants, 15.4% antipsychotics, and 7.7% antimanics.

**Table 1.** Characterization of neurotypical and 22q11.2DS individuals included in the analyses: age, sex, IQ, and working memory.

|  | Neurotypical controls | 22q11.2DS | t-test | Cohen's d |
| --- | --- | --- | --- | --- |
| Age | M=21.88; SD=6.86 | M=21.92; SD=6.89 | $t=-0.02$, $df=50.00$, $p=.98$ | $d=-0.01$ |
| Sex | 10 males, 16 females | 9 males, 17 females | - | - |
| IQ | M=112.17; SD=14.76 | M=71.40; SD=12.58 | $t=10.38$, $df=45.21$, $p<.01$ | $d=2.97$ |
| Working memory | M=103.67; SD=11.63 | M=77.16; SD=15.02 | $t=6.52$, $df=40.74$, $p<.01$ | $d=1.97$ |

Figure 1 shows the averaged ERPs and topographies for the time windows of interest (N1 and MMN), per SOA and by group. Overall, 22q11.2DS data are characterized by increased auditory evoked responses (Figure 1B) and greater variability (Figure 1D).

**Figure 1.**
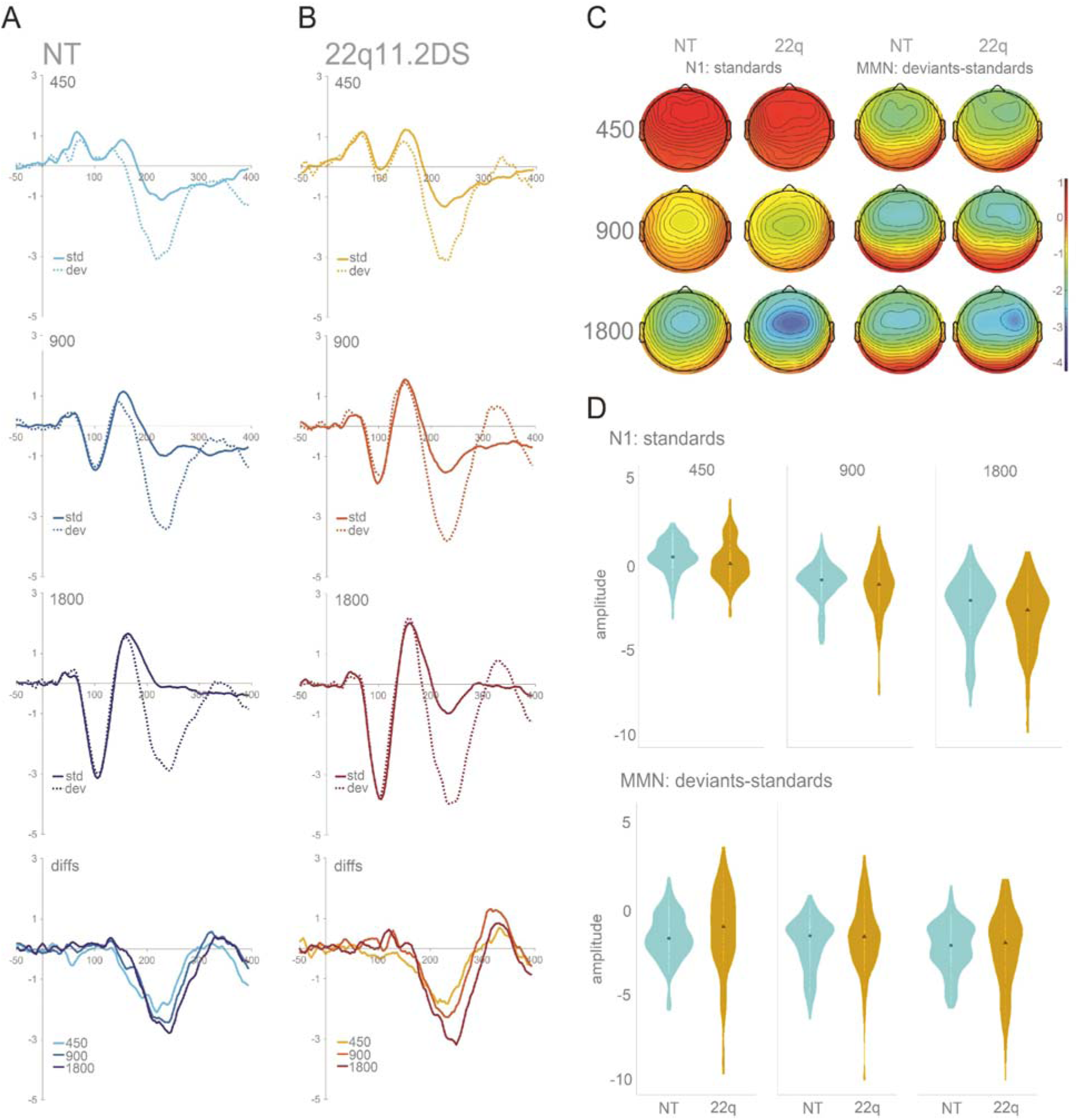
Panel A: Averaged ERPs per SOA for the NT group at FCz (fourth plot labeled as *diffs* shows MMN, i.e., differences between standards and deviants); Panel B: Averaged ERPs and topographies per SOA for the 22q11.2DS group at FCz (fourth plot labeled as *diffs* shows MMN); Panel C: Topographies for the N1 (standards only) and the MMN time windows, organized from the shorter (450 ms) to the longer (1800 ms) SOA, per group; Panel D: Violin plots showing distribution of amplitudes for N1 and MMN per group and SOA at FCz. Black square indicates median.

Separate mixed-effects models were implemented to analyze the N1 and MMN data, using the *lmer* function in the *lme4* package (87) in R (86). Mean amplitude at FCz was the numeric dependent variable. For the N1, only standards were considered. For the MMN, mean amplitude represents the difference between standards and deviants. Group (NT = −0.5, 22q11.2DS = 0.5) was a contrast-coded fixed factor, and SOA a numeric fixed factor. Subjects and SOA were added as random factors. Models were fit using the maximum likelihood criterion. *P* values were estimated using *Satterthwaite* approximations.

For the N1, while no significant effect of group was found, there was an interaction between group and SOA due to a larger N1 adaptation effect in the 22q11.2DS group when compared to the neurotypical control group. That is, the longer SOA resulted in a larger increase in N1 in the 22q group (*ß* = −0.33, SE = 0.12, *p* < .01). Still, N1 amplitude modulated as a function of SOA across both groups: Both 900 ms (*ß* = −1.54, SE = 0.06, *p* < .001) and 1800 ms (*ß* = −3.08, SE =, 0.06 *p* < .001) SOAs elicited larger responses than the 450 ms SOA. After correction for multiple comparisons, N1 amplitudes did not correlate significantly with age, IQ, working memory, or number of psychotic symptoms.

For the MMN, and as observed in the N1 time window, no significant effect of group was found, but group interacted significantly with SOA: The difference between the 450 ms and the 1800 ms SOA was again larger in the 22q11.2DS group than in the neurotypical control group (*ß* = −0.49, SE = 0.15, *p* < .001), appearing to reflect a smaller response to the shorter SOAs and a larger response to the longer SOA in the 22q11.2DS group, when compared to the neurotypical group. An effect of SOA was additionally observed: Both the 900 ms SOA (*ß* = −0.45, SE = 0.07, *p* < .001) and the 1800 ms SOA (*ß* = −0.80, SE = 0.07, *p* < .001) conditions elicited larger MMN responses than the 450 ms SOA. MMN amplitudes did not correlate significantly with age, IQ, working memory, or number of psychotic symptoms.

Given the N1 and MMN reductions described in the literature for those with schizophrenia (47, 48, 55, 56, 88) and the presence of at least one psychotic symptom in more than half of the individuals with 22q11.2DS tested here (N=15; 22q11.2DS^+^), an additional analysis was conducted in which the N1 and MMN amplitudes of those with the deletion and at least one psychotic symptom were compared to those with the deletion but no psychotic symptoms (N=11; 22q11.2DS^−^). With this analysis, we aimed at understanding whether the findings described represented the 22q11.2DS sample regardless of the presence of psychosis, or, rather, reflected the sample’s mixed nature (individuals with and without psychotic symptoms). Table 2 shows a summary of the included participants’ IQ and performance on the working memory tasks. No differences were found between the groups. In the 22q11.2DS^−^ group, 45.5% were taking antidepressants, 9.1% anticonvulsants, and 9.1% antipsychotics. In the 22q11.2DS^+^ group, 26.7% were taking antidepressants, 33.3% anticonvulsants, 20.0% antipsychotics, and 12.3% antimanics.

**Table 2.** Characterization of 22q11.2DS^−^ and 22q11.2DS^+^ individuals included in the analyses: age, sex, IQ, and working memory.

|  | 22q11.2DS<sup>-</sup> | 22q11.2DS<sup>+</sup> | t-test | Cohen's d |
| --- | --- | --- | --- | --- |
| Age | M=23.26; SD=7.75 | M=20.87; SD=6.25 | $t=0.88, df=18.78, p=.75$ | $d=0.34$ |
| Sex | 5 males, 6 females | 4 males, 11 females | - | - |
| IQ | M=73.91; SD=10.97 | M=69.43; SD=13.79 | $t=0.90, df=22.99, p=.75$ | $d=0.36$ |
| Working memory | M=81.27; SD=16.10 | M=73.93; SD=13.85 | $t=1.20, df=19.85, p=.73$ | $d=0.09$ |

Figure 2 shows the averaged ERPs and topographies for the time windows of interest (N1 and MMN), per SOA and by group (22q11.2DS^−^ and 22q11.2DS^+^).

**Figure 2.**
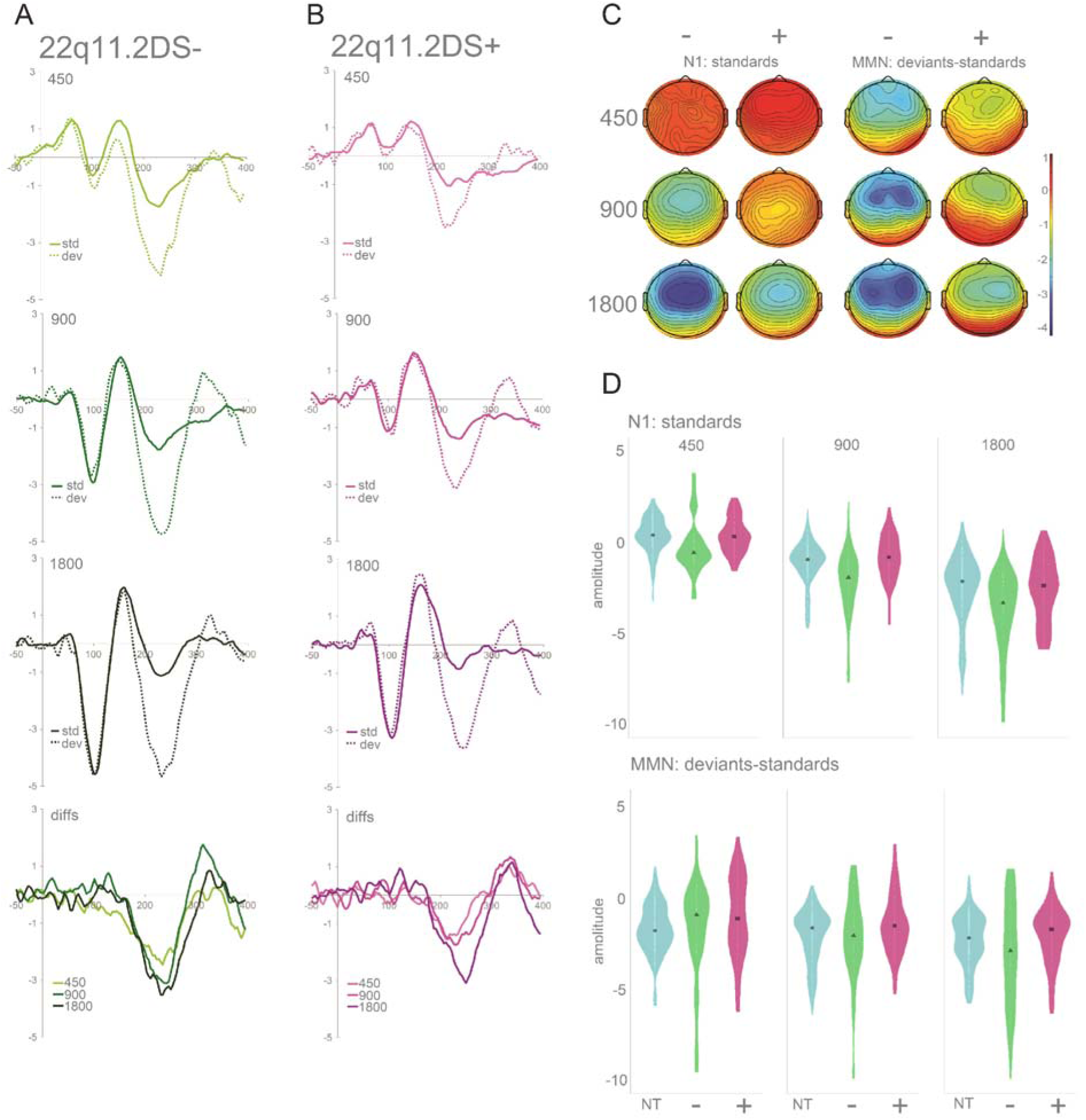
Panel A: Averaged ERPs per SOA for the group with no psychotic symptoms at FCz (fourth plot labeled as *diffs* shows MMN, i.e., differences between standards and deviants); Panel B: Averaged ERPs and topographies per SOA for the group with one or more psychotic symptoms at FCz (fourth plot labeled as *diffs* shows MMN); Panel C: Topographies for the N1 (standards only) and the MMN time windows, organized from the shorter (450 ms) to the longer (1800 ms) SOA, per group; Panel D: Violin plots showing distribution of amplitudes for N1 and MMN per group (NT group added here for comparison) and SOA at FCz. Black square indicates median.

A mixed-effects model, similar to the one described before, was implemented to test for differences among the groups. For the N1, no significant effect of group was found, but group interacted significantly with SOA, with individuals with 22q11.2DS^+^ showing reduced differences between the 450 ms and the 900 ms SOA (*ß* = 0.86, SE = 0.20, *p* < .001) and between the 450 ms and the 1800 ms SOA (*ß* = 0.62, SE = 0.20, *p* < .01) conditions, when compared to the 22q11.2DS^−^ group. As illustrated in Figure 3, when compared to neurotypical controls, the 22q11.2DS^+^ group showed the same decreased SOA effect (presenting a reduced difference between the 450 ms and the 900 ms SOA (*ß* = 0.28, SE = 0.15, *p* = .05)), whereas the 22q11.2DS^−^ group showed an increased SOA effect, with enlarged differences between the 450 ms and the 900ms SOA (*ß* = −0.58, SE = 0.16, *p* < .001) and between the 450 ms and the 1800 ms SOA (*ß* = −0.69, SE = 0.16, *p* < .01). Post-hoc analyses showed that while the 22q11.2DS^−^ group presented a significantly increased amplitude at the 900ms SOA (Tukey’s Post-hoc test, *p*<.05), the 22q11.2DS^+^ group presented a significantly decreased amplitude at the 900ms SOA (Tukey’s Post-hoc test, *p*<.01), as can be appreciated in Figure 3.

**Figure 3.**
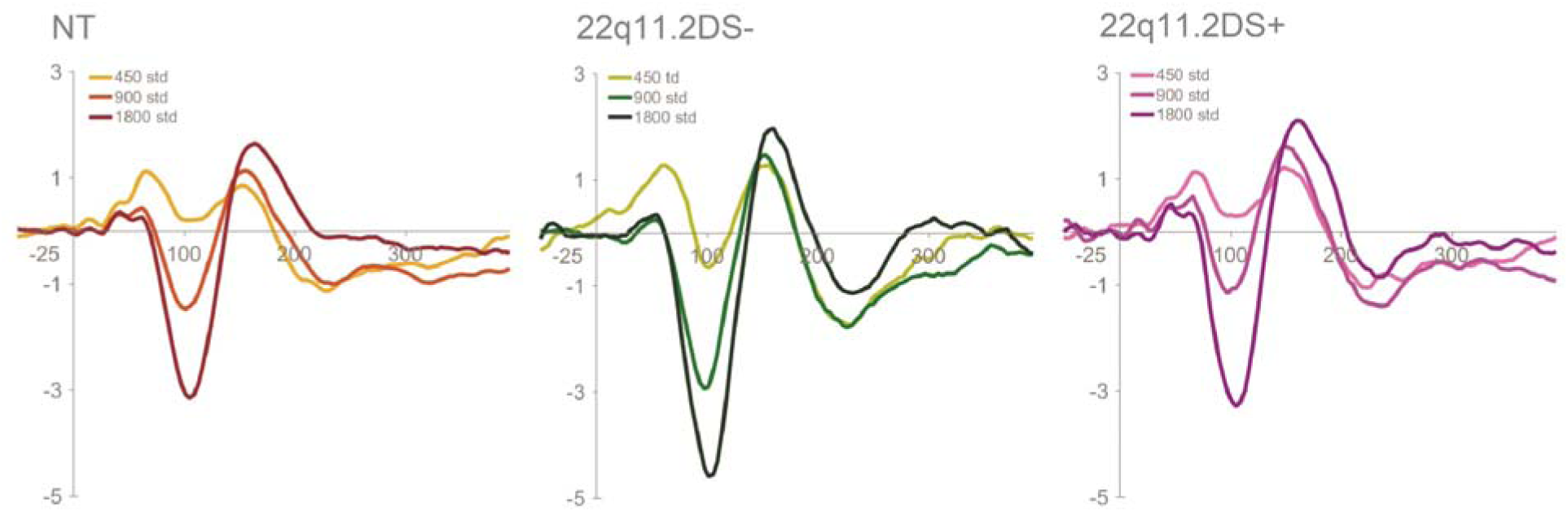
Averaged ERPs (standard tones) per group and SOA at FCz.

In the MMN time window, again, while no significant effect of group was found, a significant interaction between group and SOA was observed. Both the 22q11.2DS^+^ (*ß* = −0.46, SE = 0.17, *p* < .01) and 22q11.2DS^−^ (*ß* = −0.53, SE = 0.19, *p* < .01) groups presented an increased difference between the 450 ms and the 1800 ms SOA, when compared to neurotypical controls.

## Discussion

We characterized early auditory sensory processing using high-density EEG recordings in a group of adolescents and adults with 22q11.2DS, with and without psychotic symptomatology, and related these measures to cognitive and clinical characteristics to assess their potential informativeness with regard to vulnerability, conversion, and progression of psychosis.

First, when compared to their neurotypical peers, individuals with 22q11.2DS presented increased adaptation effects, i.e. larger N1 amplitude differences between fast and slow SOAs. The N1 response was diminished for both groups at the fastest presentation rate, but larger for the 22q11.2DS group at the slowest presentation rate. Increased auditory evoked potentials have been described in a 22q11.2DS mouse model: Loudness dependent amplitudes were enlarged in mice with the deletion (89). Likewise, two human studies looking at auditory processing in 22q11.2DS showed enhanced responses (76, 90). While larger N1s have been associated with elevated activity in the anterior cingulate and dorsomedial frontal cortex (76) and alterations in the cortical glutamate N-methyl-D-aspartate (NMDA) receptors (90–92), the nature of the neural mechanisms underlying adaptation are not fully understood. Decreases in amplitude with faster presentation rates may be observed due to temporal limitations intrinsic to the mechanisms underlying N1 generation, i.e., faster presentations of auditory stimuli do not allow for full recovery of such mechanisms and a decline in N1 amplitude is observed (43). Given that the groups showed strong adaptation for the fastest presentation rate, adaptation processes appear to be operational in 22q11.2DS. Rather, it was the responses at the slower presentation rates that differed in the 22q11.2DS group, being larger in the 22q11.2DS group overall (and in the group without symptoms). This could represent either a coarser representation of slower presentation rates, or, alternatively, a higher N1 sensory response ceiling in 22q11.2DS compared to the neurotypical population. Future work in which a larger range of SOAs is used will help to disentangle these possibilities.

Interestingly, here, we also found that N1 effects in 22q11.2DS were influenced by the presence or absence of psychotic symptoms. While the group without symptoms recapitulated what was described for the 22q11.2DS sample as a whole, individuals with one or more symptoms showed decreased differences between SOAs compared to the no symptoms group and to the neurotypical control group. This was due to a decreased N1 for the 900ms condition in the 22q11.2DS+ group (Figure 3). Decreased N1 amplitudes in the presence of psychotic symptomatology accord with findings in the schizophrenia literature, where N1 is typically reduced (45–48, 93, 94). Such a reduction has been thought of as indexing genetic risk for schizophrenia, given that reduced N1 responses were found in first-degree relatives of individuals diagnosed with schizophrenia (48). Remarkably, that does not seem to be the case in 22q11.2DS, given that those with the deletion and no psychotic symptoms (but still considered at-risk for psychosis due to the mutation) presented, instead, increased amplitudes. This could be explained by the action of two opposing mechanisms: one related to the deletion resulting in increased early sensory responses; another associated with the presence of psychotic symptomatology, which has as its outcome decreased sensory responses. Upon examination of Figure 3, one could, alternatively, argue that the 22q11.2DS+ group presents a seemingly typical N1 response function. This is, in our opinion, unlikely. Instead, we suggest that the typical looking response may be due to the two opposing mechanisms influencing the N1 response in this group. As a final point, it is also possible that N1 adaptation differences between those with and without symptoms could reflect medication effects, given that more medication intake was reported in the 22q11.2DS group with psychotic symptoms. A supplementary analysis looking at differences between those with psychotic symptoms taking one or more drugs (N=7) and those with symptoms but taking no drugs (N=8) are not suggestive of differences in amplitude between the groups, but considerably larger samples would be needed to draw confident conclusions.

Second, as described for the N1, individuals with 22q11.2DS presented increased MMN amplitude differences for fast and slow SOA conditions. Contrary to what was found for the N1, this enhancement was observed regardless of the presence of psychotic symptoms, though more apparent in the 22q11.2+ group. Previous studies on MMN in 22q11.2DS have yielded somewhat inconsistent evidence (for a review, see (95)): Whereas some studies have found reduced pitch, duration and frequency MMNs (90, 96), others failed to show differences between individuals with 22q11.2DS and their neurotypical peers (75, 97). Such inconsistencies in MMN differences may reflect the phenotypic heterogeneity that is characteristic of 22q11.2DS, as well as, possibly, methodological and stimulus related differences (e.g., (96) only report differences in frontal channels; (90) tested children and adolescents between the age of 8 and 20 years old). Adding to the range of findings, we observed that at the fast stimulation rate, which is similar to the rate often used in MMN studies (see, for instance, Cantonas et al., 2019), the response was intact and appeared typical. It was only at the longest SOA that an increase in MMN amplitude was observed relative to the MMN evoked for the shortest SOA condition. This pattern contrasts with an expected attenuation of MMN amplitude with increasing SOAs in this group, due to weaker memory traces (see (78)). Previous work from our lab using this same MMN paradigm showed that increasing SOA effectively indexed weakness in maintenance of the memory trace in Rett Syndrome (77). Given that memory deficits have been described in 22q11.2DS, we expected attenuated MMN amplitudes with longer SOAs. The current findings are thus hard to reconcile with the extant literature and, to our knowledge, have not been previously reported. They add, nevertheless, to evidence that enhanced sensory neural responses are seen in 22q11.2DS, and suggest that they are more apparent at slower presentation rates. That, contrary to what was observed for the N1, no differences between those with and without psychotic symptoms were seen for the MMN, might be indicative of the respective sensitivity of these components to the presence of psychotic symptoms. Indeed, while the matter remains one of debate, it has been suggested that the MMN deficit seen in schizophrenia is most likely reflecting disorder chronicity (70). N1 may thus represent a better endophenotype of psychosis.

Lastly, individuals with 22q11.2DS had lower IQ scores and impaired working memory, as consistently described for this population (18, 98–100). Though intellectual disability is quite prevalent in 22q11.2DS, its causes are not well understood. Several genes in the deleted region have been identified as potential candidates: *Tbx1* has been implicated in brain development and may thus have a role in cognitive deficits (101); PRODH mutations are associated with intellectual disability, and are found in about one-third of the individuals diagnosed with 22q11.2DS (102). Proline, the enzyme produced by PRODH, in interaction with catechol-O-methyl-transferase (COMT) seems to negatively impact brain function in children with 22q11.2DS (103, 104). Regarding working memory, the copy number elevations of COMT have been associated with such impairments, not only in 22q11.2DS (105), but also in schizophrenia (106) (but see (107) for different findings). COMT mutations have been argued to result in increased dopamine degradation in the frontal lobes, which could provide a molecular basis for some of the symptomatology associated with both schizophrenia and 22q11.2DS (108, 109). Given that cognitive impairment is characteristic of 22q11.2DS, it is important to note that, while the cognitive decline that seems to precede and predict the development of psychosis in idiopathic schizophrenia (110, 111) might be somewhat conspicuous, such a decline may not be as noticeable in 22q11.2DS and might require more thorough monitoring. Indeed, in 22q11.2DS, cognitive deficits may be traits that preexist and increase risk for psychosis, but, keeping in mind this shifted baseline, should still allow one to discriminate those most susceptible to psychotic symptoms (112). Additionally, increased variability was observed in the 22q11.2DS group, both in the N1 and the MMN time windows, in line with the remarkable variability in expression described in the syndrome (6, 113–115). This observation concords with the presence of subgroups within 22q11.2DS characterized by different cognitive and neural profiles, and with different vulnerability for schizophrenia. Though here we focus on the distinction between individuals with and without psychotic symptoms, other subgroups with different cognitive and neural profiles may co-exist within 22q11.2DS. The definition of such subgroups would not only add to the understanding of the phenotype, but also, potentially, impact intervention.

Some limitations to this study should be noted. Despite the substantial size of our sample considering the rare nature of 22q11.2DS, larger numbers would allow for more detailed analyses (particularly those looking at associations between neural, cognitive and behavioral outcomes) and to take into consideration the impact of medication on the findings reported. In future work, different measures of psychotic symptoms should be used to better characterize psychosis in this population. Instead of or in addition to the diagnostic interview used here, measures providing symptom severity and the differentiation between negative and positive symptoms could be more informative.

## Acknowledgements

We wish to thank Drs. Juliana Bates, Katherine Behar, and Pamela Counts who performed the clinical assessments, and Elise Taverna and Danielle Newbury for their help with data collection. We would also like to acknowledge the role of the Montefiore-Einstein Regional Center for 22q11.2 Deletion Syndrome in recruitment, and the Rose F. Kennedy Intellectual and Developmental Disability Research Center for all of its support. We extend our most sincere gratitude to the participants and their families for their interest, their involvement, and their time. This work was supported in part by the Eunice Kennedy Shriver National Institute of Child Health and Human Development (NICHD), under award number U54 HD090260.

## Disclosures

The Authors report no biomedical financial interests or potential conflicts of interest.

